# 18S rRNA 3’ end cleavage by the phosphorylated endoribonuclease NOB1 is interconnected with early small subunit biogenesis

**DOI:** 10.64898/2026.06.29.735188

**Authors:** Katja I. Bloch von Blottnitz, Mona Honemann-Capito, Philipp Hackert, Olexandr Dybkov, Christof Lenz, Markus T. Bohnsack, Sonja Lorenz, Henning Urlaub, Claudia Schneider, Katherine E. Bohnsack

## Abstract

Processing of the precursor ribosomal RNAs (pre-rRNAs) is a key aspect of ribosomal subunit assembly that is closely coordinated with other maturation events. The ribonucleases that mediate pre-rRNA cleavages require regulation to ensure that their activities are exerted in a timely manner. Post-translational modifications can influence protein functions, and although many human ribosome assembly factors are reported to be post-translationally modified, most of these sites remain unconfirmed and functional insights are lacking. Here, we show that NOB1, the PIN domain endoribonuclease responsible for cleavage of the 3′ end of the 18S rRNA, is phosphorylated within an evolutionarily conserved acidic tract that can be modified by casein kinase II *in vitro*. Association of NOB1 with pre-ribosomes is independent of these phosphorylations, and lack of NOB1 phosphorylation only mildly perturbs the efficiency of SSU maturation events upstream of 3′ end cleavage of the 18S rRNA. Interestingly, our analyses of pre-rRNA levels in cells depleted of NOB1 or lacking its catalytic activity revealed not only accumulation of the 18SE precursor of the 18S rRNA, but also altered levels of pre-rRNAs containing 5′ external transcribed spacer (ETS) sequences (43S, 26S and 30S). This suggests that lack of NOB1-mediated pre-rRNA cleavage impairs recycling of assembly factors required during early biogenesis steps, leading to altered kinetics of 5′ ETS processing. Taken together these data provide new insights into the role of NOB1 during SSU biogenesis and the post-translational regulation of this ribonuclease.

## INTRODUCTION

Ribosomes are multi-megadalton ribonucleoprotein complexes responsible for protein synthesis in cells. Eukaryotic cytosolic ribosomes are composed of four ribosomal RNAs (rRNAs; 18S, 5.8S, 28S and 5S) and 80 ribosomal proteins (RPs) arranged into a small (SSU; 40S) and large subunit (LSU; 60S) (Vanden Broeck and Klinge 2024; Bohnsack and Bohnsack 2019). The SSU contains the decoding centre, whereas the LSU harbours the peptidyl transferase centre and the polypeptide exit tunnel (Khatter et al. 2015). Assembly of the ribosomal subunits is one of the most energetically costly cellular processes and requires the coordinated action of all three RNA polymerases (Pol) (Warner 1999). Pol II transcribes *RP* genes and Pol III synthesises the 5S rRNA whereas Pol I-mediated rDNA transcription leads to production of a pre-rRNA transcript (47S) containing the mature 18S, 5.8S and 28S rRNAs interspersed by internal transcribed spacers (ITS1 and ITS2) and flanked by external transcribed spacers (5′ ETS and 3′ ETS) (Henras et al. 2015). Assembly of the RPs onto the rRNA scaffold is facilitated by >200 ribosome assembly factors (AFs) that include enzymes responsible for the modification, folding and processing of the pre-rRNAs as well as proteins that scaffold the assembly process (Aubert et al. 2018; Dörner et al. 2023; Sloan et al. 2017). Due to the high demand for ribosomes in proliferating cells, recycling of *trans*-acting AFs needs to be effective and malfunctions thereof can lead to defects in the assembly pathway.

Processing of the 47S pre-rRNA is initiated co-transcriptionally by cleavage at the A′ site in the 5′ ETS (Lazdins et al. 1997). Subsequent endoribonucleolytic cleavages in the 5′ ETS at sites A0 and 1, likely performed by UTP23 and UTP24, respectively, generate the mature 5′ end of the 18S rRNA (Tomecki et al. 2015; Wells et al. 2016, 2017). The spacer fragments excised by these sequential endoribonucleolytic cleavages are exoribonucleolytically degraded by XRN2 and the RNA exosome within the context of the SSU processome, the first stable pre-ribosomal intermediate (Singh et al. 2021; Sloan et al. 2014). Coordinated with cleavages at site A0 and 1, processing at site 2 in ITS1 by the ribozyme RNase MRP separates the pre-rRNAs of the SSU and LSU (Goldfarb and Cech 2017). For maturation of the 18S rRNA, cleavage at site 2 is typically followed by 3′-5′ exoribonucleolytic trimming by the EXOSC10-containing RNA exosome and ISG20L2 towards site E (also known as 2a) to generate the 18SE pre-rRNA; however, site E can also be cleaved directly by UTP24 to form 18SE (Aubert et al. 2025; Preti et al. 2013; Sloan et al. 2013; Wells et al. 2016). Site E cleavage is followed by further 3′-5′ trimming by the polyA-specific ribonuclease PARN (Montellese et al. 2017). Formation of the mature 3′ end of the 18S rRNA is achieved via cleavage at site 3 by the endoribonuclease NOB1 (O’Donohue et al. 2010; Sloan et al. 2019).

NOB1 is a PilT N-terminal (PIN) domain endoribonuclease containing four conserved acidic residues (aspartate (D)10, glutamate (E)36, D86 and D238) in its active site (Pertschy et al. 2009; Veith et al. 2012; Raoelijaona et al. 2018). Also present in NOB1 is a zinc finger domain (ZNF) that mediates interactions with helix (h)36 in the 3′ major domain of the 18S rRNA (Veith et al. 2012). NOB1 associates with pre-40S particles in the nucleus, but is initially held at a distance from its substrate, the 18S–ITS1 junction (site 3) (Ameismeier et al. 2018). Within nuclear pre-SSU particles, the C-terminal region of NOB1 and the KH domains of PNO1 mask site 3, and these two proteins maintain helices (h)44/45 of the 18S pre-rRNA in an immature conformation. Following export of pre-SSU particles to the cytosol, various co-exported pre-SSU AFs are released and the pre-rRNAs assume their mature conformations (Cerezo et al. 2019). Detachment of TSR1 and LTV1 triggers final positioning of h44/45, which in turn promotes recruitment of RIOK1 and dissociation of PNO1. Release of PNO1 induces a substantial conformational rearrangement in NOB1 that re-positions the enzyme such that site 3 is placed within its catalytic site and cleavage of the 3′ end of the 18S rRNA rapidly follows (Plassart et al. 2021; Ameismeier et al. 2020). The catalytic activity of NOB1 in pre-SSU complexes is thus regulated by structural constraints, ensuring that its active site only contacts its substrate at the appropriate stage (Ameismeier et al. 2018, 2020; Plassart et al. 2021). Following site 3 cleavage, NOB1 then dissociates, likely together with the cleaved fragment of ITS1 (Parker et al. 2019; Ameismeier et al. 2020).

As pre-rRNA cleavages are committed steps during ribosomal subunit assembly, regulation of the ribonucleases involved is critical (Schneider and Bohnsack 2023). Several kinase signalling pathways are linked to ribosome biogenesis and phosphorylation marks on various RPs and AFs have been identified (reviewed in (Lawrence et al. 2026)). For example the TORC1 kinase phosphorylates conserved sites within the RP eS6 (Meyuhas 2015), and yeast Tor1 and casein kinase II control a switch between productive and non-productive pre-rRNA processing pathways (Kos-Braun et al. 2017). During SSU biogenesis, casein kinase I phosphorylates the AFs LTV1 and ENP1 as well as the RP eS3, promoting stable incorporation of eS3 into nuclear pre-SSU and their subsequent export to the cytosol (Zemp et al. 2014). Furthermore, RSK phosphorylates RIOK2, triggering its release and the dissociation of other AFs from maturing pre-SSU particles (Cerezo et al. 2021). RIOK2 itself was shown to phosphorylate another pre-SSU AF, DIMT1L *in vitro* (Sloan et al. 2019). Many other AFs have been identified in high-throughput screens for phosphorylated proteins; however, most putative sites lack robust experimental confirmation and it remains unknown whether these marks influence protein functions (Needham et al. 2019; Hornbeck et al. 2015). Phosphoproteomics indicate that NOB1 may be phosphorylated (Hornbeck et al. 2015) and here, we map modification sites and investigate their influence on NOB1 function in SSU biogenesis. Interestingly, it was further recognised in the course of these analyses that 18S 3′ end maturation by NOB1 is coupled to efficient processing of the 5′ ETS, likely through the recycling of AFs.

## RESULTS AND DISCUSSION

### Human NOB1 is phosphorylated at specific sites

To ascertain whether NOB1 is a phosphoprotein, as suggested by high-throughput analyses (Hornbeck et al. 2015), cell extracts were prepared in the presence of phosphatase inhibitors (PI) to retain endogenous phosphorylations or treated with alkaline phosphatase (AP) to non-specifically remove them. Upon separation of the extracts in denaturing Bis-Tris gels, NOB1 was detected as a single band, migrating similarly irrespective of the treatment (**Fig. 1A**). By contrast, separation of the cell extracts in gels supplemented with the ‘Phos-tag’ reagent, which retards the migration of phosphorylated proteins, revealed a shift in NOB1 migration between the two extracts, demonstrating that NOB1 is indeed a phosphoprotein. Note that dephosphorylation by alkaline phosphatase i) sometimes reduced the level of NOB1 and ii) was not always complete, therefore resulting, at times, in partially dephosphorylated states.

**Figure 1.**
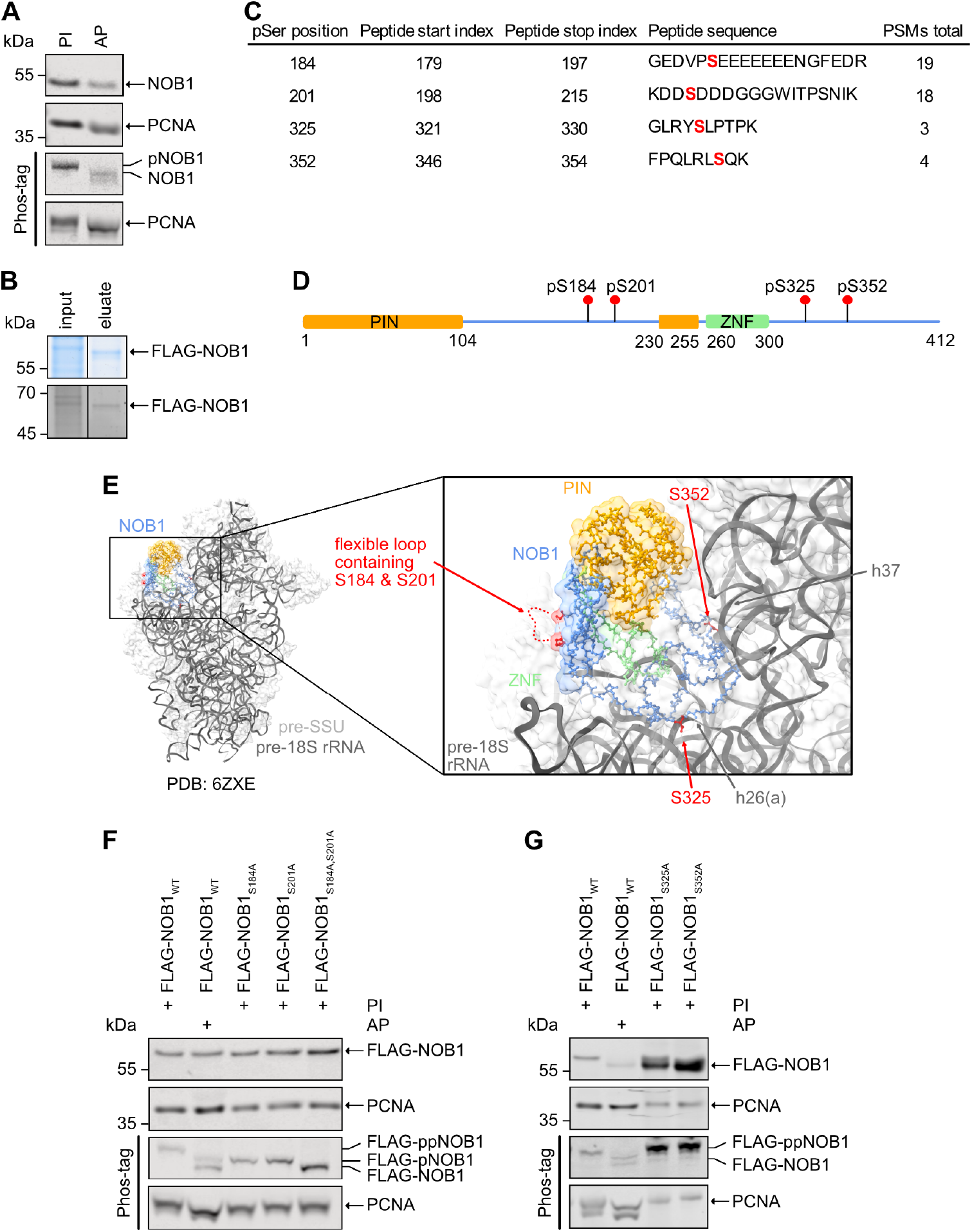
Mapping NOB1 phosphorylation sites. **(A)** Extracts from HeLa cells were prepared in the presence of phosphatase inhibitors (PI) or were treated with alkaline phosphatase (AP). Proteins in the extracts were separated by electrophoresis on Bis-Tris gels supplemented or not with Phos-tag reagent, and the endogenous NOB1 was detected by western blotting. PCNA, which is itself phosphorylated, served as a loading control. pNOB1 indicates phosphorylated NOB1. **(B)** A phosphatase inhibitor-treated extract from cells expressing FLAG-NOB1 was used for anti-FLAG IP, and proteins in the input (0.2%) and eluate were detected by Coomassie staining (upper) and phosphoprotein gel staining (lower). **(C)** Samples prepared as in (B) were subjected to phosphorylation mass spectrometry and the table shows the identified phosphorylated residues in NOB1 and their metrics. PSMs refers to peptide-spectrum matches. **(D)** Scheme showing the domain architecture of NOB1 with phosphoserines (pS) identified by phosphorylation mass spectrometry highlighted. Numbers below the scheme indicate amino acids at domain boundaries according to (Raoelijaona et al. 2018). PilT N-terminal domain – PIN (orange); zinc finger – ZNF (green). **(E)** Visualisation of the amino acids identified as phosphorylated in (C) on the structure of NOB1 (blue; PIN domain – orange; ZNF – green) within a selected pre-SSU complex (PDB: 6ZXE). 18S rRNA helices (h) proximal to NOB1 phosphorylation sites are indicated. Note that S184 and S201 are in a flexible region not resolved in any available pre-SSU structure (indicated by a dotted line), so the closest resolved amino acids are highlighted. **(F)** Extracts from cells expressing FLAG-NOB1_WT_ or FLAG-NOB1 with serine to alanine substitutions at the positions 184, 201 or both were prepared in the presence of phosphatase inhibitors (PI) or were treated with alkaline phosphatase (AP) as indicated. Proteins in the extracts were separated by electrophoresis on Bis-Tris gels supplemented or not with Phos-tag reagent, and the indicated proteins were detected by western blotting (anti-FLAG and anti-PCNA). pNOB1 and ppNOB1 indicate NOB1 with one or two detectable phosphorylations, respectively. **(G)** Experiments were performed as in (F) using cells transiently expressing FLAG-NOB1_WT_ or FLAG-NOB1 with serine to alanine substitutions at the positions 325 or 352. All experiments were performed in triplicate and representative data are shown.

To identify NOB1 phosphorylation sites in HeLa Flp-In cells, a stably transfected cell line for inducible expression of FLAG-NOB1 was generated (**Supplementary Fig. S1A**), and FLAG-NOB1 was enriched by anti-FLAG IP under stringent conditions in the presence of phosphatase inhibitors. Following confirmation of NOB1 phosphorylation by phosphoprotein gel staining (**Fig. 1B**), the band corresponding to FLAG-NOB1 was subjected to phosphorylation mass spectrometry (MS), which identified serines (S)184, 201, 325 and 352 as phosphorylation sites (**Fig. 1C**). Although exact modification stoichiometries cannot be obtained using this approach, the presence of phosphoserine (pS) at positions 184 and 201 was detected more robustly than those at positions 325 and 352. S184 and S201 reside in a long linker region within the split NOB1 PIN domain (Raoelijaona et al. 2018), whereas S325 and S352 are present downstream of the zinc finger (ZNF) that is required for RNA binding (**Fig. 1D**) (Sloan et al. 2019). Within all the available structures of pre-SSU particles, S325 and S352 are orientated towards helices 26(a) and 37 of the 18S rRNA, respectively, whereas pS184 and pS201 are in a flexible loop that is not resolved (**Fig. 1E**).

Cells expressing FLAG-tagged versions of NOB1 containing serine to alanine (A) substitutions at the detected phosphorylation sites were generated (**Supplementary Fig. S1B**) to determine which pS(s) affect(s) NOB1 migration during Phos-tag gel electrophoresis. Compared to FLAG-NOB1_WT_ in phosphatase inhibitor-treated cell extracts, treatment with alkaline phosphatase caused a downward shift and splitting of the wild-type NOB1 band (**Fig. 1F**), as observed for endogenous NOB1 (**Fig. 1A**). Although alkaline phosphatase treatment sometimes reduced the level of FLAG-NOB1_WT_, the migration profile of this sample can serve as a marker for partially/non-phosphorylated protein. Phospho-null substitutions at both S184 and S201 of FLAG-NOB1 caused similar migration to fully dephosphorylated FLAG-NOB1_WT_ (FLAG-NOB1_WT_ + AP; lower band), whereas individual substitutions of either S184 or S201 led to migration similar to the partially dephosphorylated, alkaline phosphatase-treated FLAG-NOB1_WT_ sample (FLAG-NOB1_WT_ + AP; upper band) (**Fig. 1F**). Substitution of either S325 or S352 for alanine did not influence the migration profile of FLAG-NOB1 (**Fig. 1G**), consistent with the MS-based indication that these residues are phosphorylated at low levels. As the presence of serines at these or equivalent positions is conserved from humans to yeast (S325) and across metazoa (S352) (Raoelijaona et al. 2018), it is possible that these post-translational modifications are dynamically regulated and become functionally relevant only in specific conditions. Therefore, in this context, further analyses were focused on the constitutively phosphorylated S184 and S201.

### NOB1 phosphorylation within an evolutionarily conserved acidic region does not affect pre-ribosome association

Closer inspection of the context of pS184 and pS201 revealed that they lie within an acidic region immediately upstream of the PNO1 interaction motif (^208^WIT^210^) (Raoelijaona et al. 2018), and alignment of NOB1 orthologues highlights that this region is highly conserved across eukaryotes (**Fig. 2A**). Strikingly, in fly, zebrafish, worm and yeast, the residues homologous to pS184 and pS201 in human (*Hs*)NOB1 are aspartate or glutamate, indicating the conservation of acidic residues at these positions (**Fig. 2A**). Similarly, in *Mus musculus* (*Mm*; mouse), S184 is replaced by glutamine (E181), whereas S201 is conserved (S192). To determine whether S192 in *Mm*NOB1 is also phosphorylated, FLAG-tagged wild-type *Mm*NOB1 and a version containing an S192A substitution were expressed in mouse C2C12 cells. Phos-tag gel analysis showed that alkaline phosphatase treatment of the extract from cells expressing the wild-type protein led to a faster migrating version of FLAG-*Mm*NOB1, indicating partial dephosphorylation and that *Mm*NOB1 is also a phosphoprotein (**Fig. 2B**). FLAG-*Mm*NOB1_S192A_ migrated similarly to the dephosphorylated population of FLAG-*Mm*NOB1 (**Fig. 2B**), demonstrating that, like S201 of *Hs*NOB1, S192 of *Mm*NOB1 is phosphorylated. Serine/threonine kinases typically recognise their substrates via sequence motifs and a subset of kinases display preferences for phosphorylating within acidic sequence contexts (Kemp and Pearson 1990; Johnson et al. 2023). High-throughput screening of kinase recognition motifs and correlation of the identified motifs with phosphorylation sites detected in human proteins allowed prediction of kinases potentially responsible for phosphorylation of S184 and S201 of NOB1 (Johnson et al. 2023). Among the top ten candidates for pS184 and pS201, some of which could be excluded based on their cell type-specific expression profiles (**Supplementary Table S1**) (Karlsson et al. 2021; Uhlén et al. 2015), was casein kinase II (CK2), an enzyme previously linked to ribosomal subunit biogenesis (Kubiński and Masłyk 2017; Kos-Braun et al. 2017; Thomé et al. 2026; Vanden Broeck and Klinge 2023).

**Figure 2.**
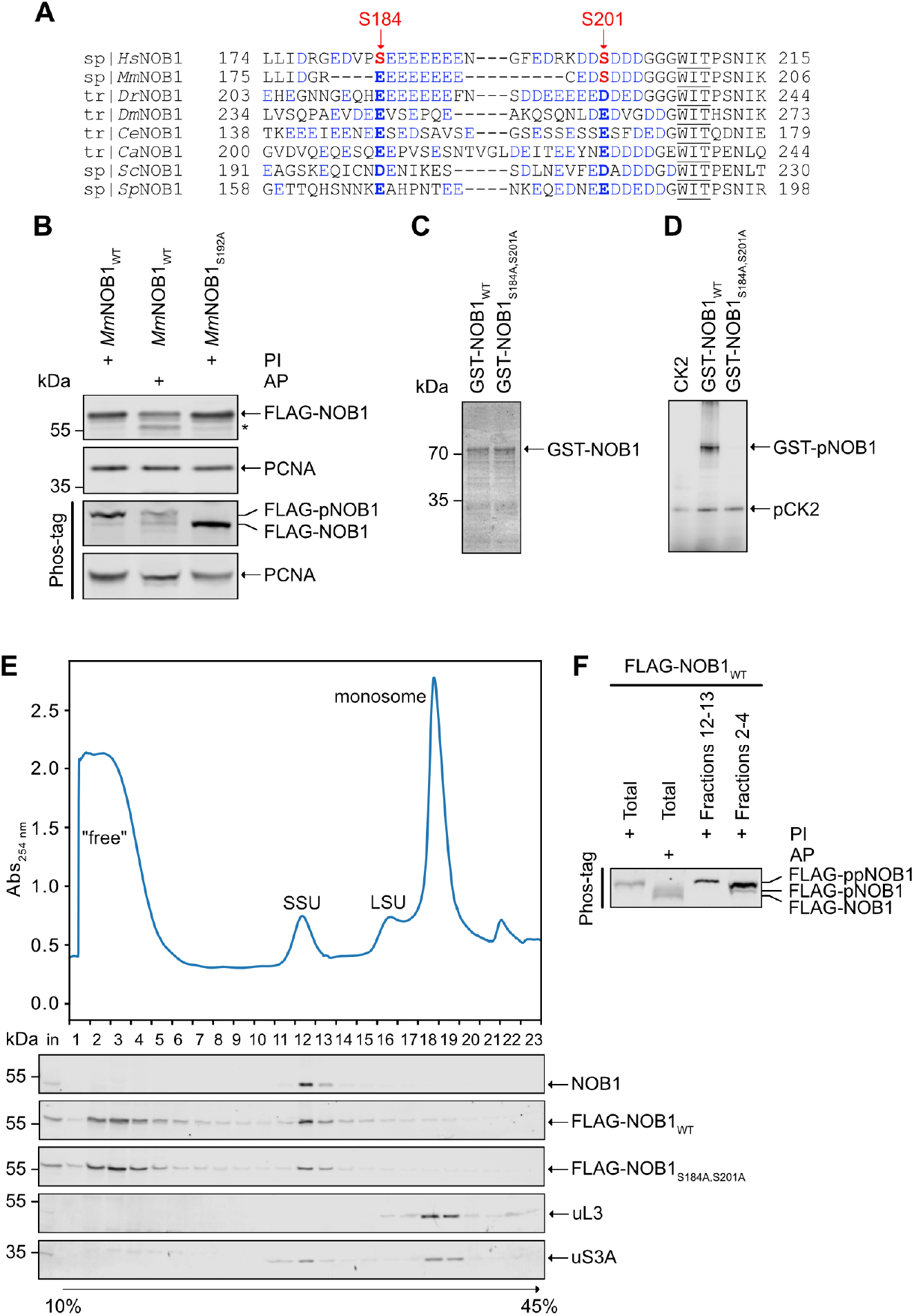
Context and conservation of S184 and S201, and pre-ribosome association of phosphorylated/phospho-null NOB1. **(A)** NOB1 protein sequences from the indicated species were aligned and the region from amino acid 174 to 215 of human (*Hs*) NOB1 is shown. Abbreviations: *Mm* – *Mus musculus, Dr* – *Danio rerio, Dm* – *Drosophila melanogaster, Ce* – *Caenorhabditis elegans, Ca* – *Candida albicans, Sc* – *Saccharomyces cerevisiae, Sp* – *Schizosaccharomyces pombe*. Acidic residues are shown in blue, human NOB1 phosphorylation sites are shown in red and equivalent positions in other homologies are highlighted in bold. The WIT motif via which NOB1 interacts with PNO1 is underlined. **(B)** Extracts from mouse C2C12 cells transiently expressing FLAG-*Mm*NOB1 or FLAG-*Mm*NOB1 with a serine to alanine substitution at position 192 (*Mm*NOB1_S192A_; equivalent to *Hs*NOB1_S201A_) were prepared in the presence of phosphatase inhibitors (PI) or were treated with alkaline phosphatase (AP) as indicated. Proteins in the extracts were separated by electrophoresis on Bis-Tris gels supplemented or not with Phos-tag reagent, and the indicated proteins were detected by western blotting (anti-FLAG and anti-PCNA). * partial degradation product. **(C)** Recombinant GST-NOB1_WT_ and GST-NOB1_S184A,S201A_ expressed in *E. coli* were purified, and are visualised following denaturing Bis-Tris gel electrophoresis and Coomassie staining. **(D)** An *in vitro* phosphorylation assay was performed using equal amounts of purified recombinant GST-NOB1_WT_ or GST-NOB1_S184A,S201A_ together with casein kinase II (CK2) as well as casein kinase II alone in the presence of γ[^32^P]-labelled ATP. Reaction products were separated by denaturing Bis-Tris gel electrophoresis and visualised using a phosphorimager. **(E)** Phosphatase inhibitor-treated extracts from HeLa cells and HeLa cell lines expressing FLAG-NOB1_WT_ or FLAG-NOB1_S184A,S201A_ were separated by sucrose density gradients. The absorbance profile of the FLAG-NOB1_WT_ gradient at 254 nm is shown with peaks corresponding to (pre-)SSU, (pre-)LSU and monosomes/90S pre-ribosomes indicated. Equivalent profiles were recorded for all gradients (see **Supplementary Fig. S2**). Proteins in each fraction of the three gradients were analysed by western blotting using antibodies against NOB1 (upper), the FLAG tag (FLAG-NOB1_WT_ and FLAG-NOB1_S184A,S201A_) or the ribosomal proteins uL3 or uS3A (lower). The western blots for uL3 and uS3a presented here derive from the FLAG-NOB1_WT_ gradient (see **Supplementary Fig. S2** for other gradients). **(F)** Fractions 2-4 and 12-13 from the FLAG-NOB1_WT_ gradients (E) containing non-pre-ribosome-associated NOB1 and pre-SSU particles, respectively, were separately pooled and the phosphorylation status of FLAG-NOB1_WT_ was analysed by Phos-tag gel electrophoresis followed by western blotting using an antibody against the FLAG tag. All experiments were performed in duplicate (E and F) or triplicate (B-D) and representative data are shown.

To interrogate CK2 activity towards NOB1 *in vitro*, GST-NOB1_WT_ and GST-NOB1_S184A,S201A_ were recombinantly expressed in *Escherichia coli* and purified (**Fig. 2C**). GST-NOB1_WT_ or GST-NOB1_S184A,S201A_ were incubated with recombinant CK2 and γ[^32^P]-labelled ATP, and protein phosphorylation was monitored autoradiographically. Alongside CK2 autophosphorylation (Pagano et al. 2005; Donella-Deana et al. 2001), phosphorylation of wild-type NOB1, but not NOB1_S184A,S201A_ (**Fig. 2D**) was detected, indicating that CK2 can phosphorylate S184 and/or S201 of NOB1.

A possible influence of NOB1 phosphorylation on its association with pre-ribosomes was then investigated. Extracts from HeLa cells and those expressing FLAG-NOB1_WT_ or phospho-null FLAG-NOB1_S184A,S201A_ were separated on sucrose density gradients. Comparison to the absorbance profile at 254 nm as well as the migration profiles of the RPs uL3 and uS3A as determined by western blotting, indicated that endogenous NOB1 migrates exclusively in fractions containing pre-SSU complexes (fractions 12-13; **Fig. 2E; Supplementary Fig. S2**). FLAG-NOB1_WT_ was enriched with pre-SSU complexes as well, but a portion was also detected in fractions containing free proteins and low molecular weight complexes (**Fig. 2E; Supplementary Fig. S2**). As this was not observed for the endogenous protein in the absence of FLAG-NOB1 expression, the non-pre-ribosome-associated pool of FLAG-NOB1 likely arises due to overexpression of the FLAG-tagged protein (**Supplementary Fig. S1A**). The fractions containing pre-SSU complexes (12-13), as well as those containing free overexpressed FLAG-NOB1_WT_ (2-4) were pooled and the phosphorylation status of FLAG-NOB1_WT_ was analysed by Phos-tag gel electrophoresis. This confirmed that both pre-ribosome-associated and free NOB1 are phosphorylated, with only a very minor portion of non-pre-ribosome-associated NOB1 partially dephosphorylated (**Fig. 2F**). Comparing the distributions of FLAG-NOB1_WT_ and phospho-null FLAG-NOB1_S184A,S201A_ in sucrose density gradients (**Fig. 2E**) revealed similar migration patterns, demonstrating that lack of S184 and S201 phosphorylation does not prevent NOB1 association with pre-ribosomes.

### Lack of NOB1 or its catalytic activity affect kinetics of 5′ ETS processing whereas loss of NOB1 phosphorylation mildly perturbs ITS1 processing upstream of NOB1 cleavage

Next, to address whether pS184 and pS201 affect the function of NOB1 in pre-rRNA processing, the levels of pre-rRNA intermediates of the pre-SSU (**Fig. 3A**) were first examined in cells depleted of NOB1 by RNAi (**Fig. 3B; Supplementary Fig. S3A**). Consistent with its role as the endoribonuclease cleaving the 3′ end of the 18S rRNA and as previously observed (Sloan et al. 2013; Preti et al. 2013; Montellese et al. 2017), depletion of NOB1 caused accumulation of the 18SE pre-rRNA (**Fig. 3C, D; Supplementary Fig. S3B**). Strikingly, it also caused accumulation of the 26S and 43S pre-rRNAs, which extend from the A0 cleavage site in the 5′ ETS to site 2 in ITS1 and site 02 at the 3′ end of the 28S rRNA, respectively, as well as a reduction in the level of the 30S pre-rRNA that is generated by cleavages at A′ in the 5′ ETS and site 2 in ITS1 (**Fig. 3C, D; Supplementary Fig. S3B**). Taken together, these alterations in pre-rRNA levels suggest that the kinetics of 5′ ETS processing are altered in the absence of NOB1 such that the 30S pre-rRNA is more rapidly converted into 26S (and the 45S into 43S), which then accumulate as subsequent pre-SSU maturation steps become limiting.

**Figure 3.**
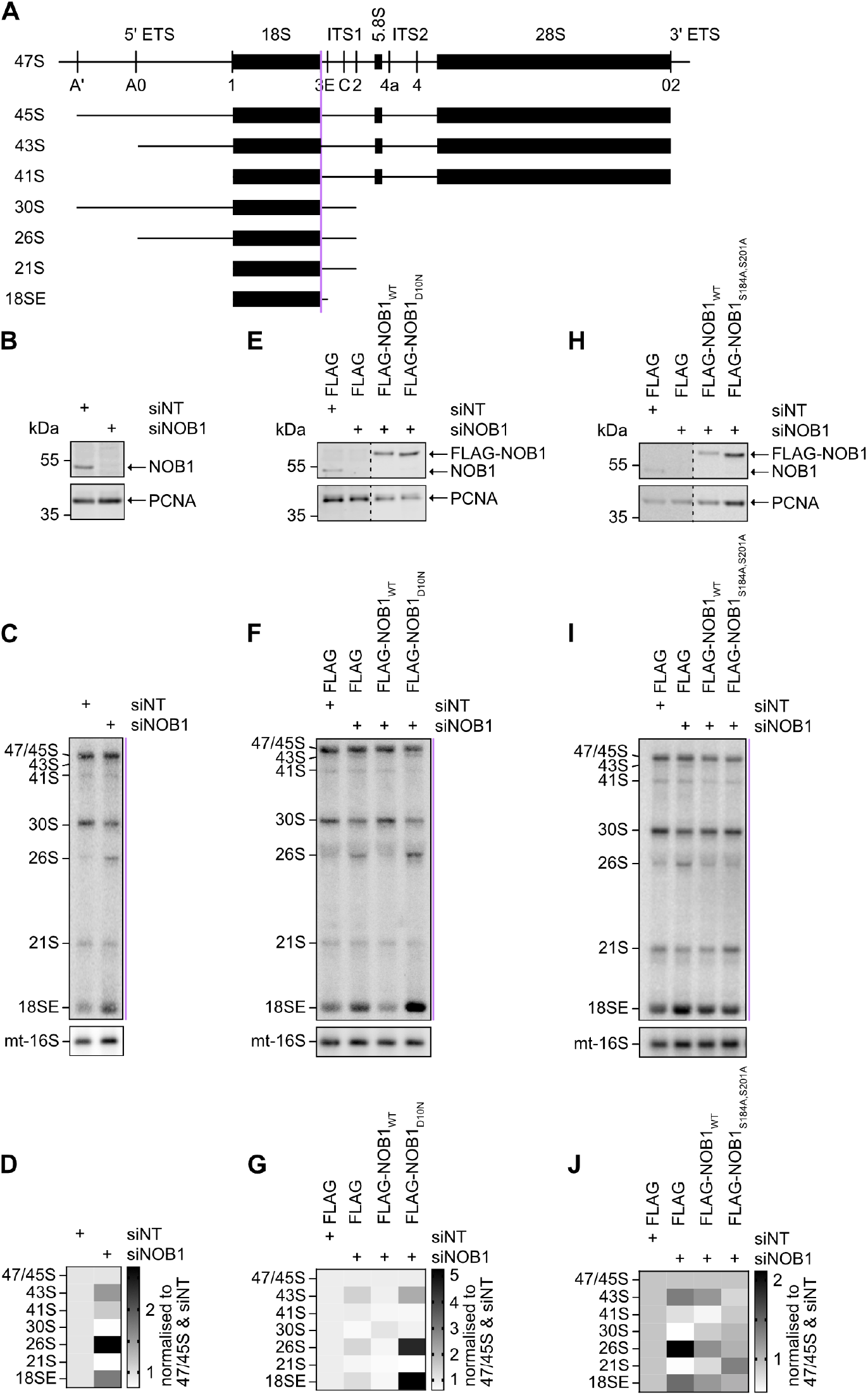
Influence of NOB1 depletion, catalytic activity and phosphorylation on pre-rRNA processing. **(A)** Scheme of the 47S pre-rRNA transcript with pre-rRNA cleavage sites marked and other pre-rRNA intermediates of the pre-SSU shown. The hybridisation position of the probe used for northern blotting is indicated by a purple line. **(B)** Proteins from HeLa cells treated with a non-target siRNA (siNT) or siRNAs targeting NOB1 (siNOB1) were analysed by western blotting using anti-NOB1 and anti-PCNA antibodies. **(C)** Pre-rRNAs from cells treated as in (B) were analysed by northern blotting using probes hybridising to the 5’ end of ITS1 and the 16S mt-rRNA. **(D)** Pre-rRNA intermediates in (C) were quantified, normalised to 47S/45S and scaled to siNT, and are presented as a heatmap (see also **Supplementary Figure S3B**). **(E)** Proteins from HeLa cells treated with siNT or siNOB1 and expressing the FLAG tag, FLAG-NOB1_WT_ or FLAG-NOB1_D10N_ were analysed by western blotting using anti-NOB1 and anti-PCNA antibodies. **(F)** Pre-rRNAs from cells treated as in (E) were analysed by northern blotting using probes hybridising to the 5’ end of ITS1 and the 16S mt-rRNA. **(G)** Pre-rRNA intermediates in (F) were quantified, normalised to 47S/45S and scaled to siNT, and are presented as a heatmap (see also **Supplementary Figure S3D**). **(H)** Proteins from HeLa cells treated with siNT or siNOB1 and expressing the FLAG tag, FLAG-NOB1_WT_ or FLAG-NOB1_S184A,S201A_ were analysed by western blotting using anti-NOB1 and anti-PCNA antibodies. **(I)** Pre-rRNAs from cells treated as in (H) were analysed by northern blotting using probes hybridising to the 5’ end of ITS1 and the 16S mt-rRNA. **(J)** Pre-rRNA intermediates in (I) were quantified, normalised to 47S/45S and scaled to siNT, and are presented as a heatmap (see also **Supplementary Figure S3F**). All experiments were performed in duplicate (H-J) or triplicate (B-G) and representative data are shown. Quantifications shown in (D, G, J) are presented as mean of the independent replicates.

To determine whether the effects of NOB1 depletion on these early pre-rRNAs arise due to lack of the NOB1 protein or its catalytic activity, a complementation system was established. In this system, endogenous NOB1 is depleted by RNAi targeting the 3′ UTR, followed by expression of FLAG-tagged wild-type NOB1 or a catalytically impaired variant in which aspartate 10 within the PIN domain is substituted for asparagine (D10N) (**Fig. 3E; Supplementary Fig. S3C**) (Sloan et al. 2019). Northern blot analysis of pre-rRNAs from these cells showed that re-expression of FLAG-NOB1_WT_ in cells depleted of endogenous NOB1 restored 43S, 30S, 26S and 18SE levels to be comparable to those in control cells (**Fig. 3F, G; Supplementary Fig. S3D**), confirming that these pre-rRNA processing defects arise due to the specific loss of NOB1. When expressing FLAG-NOB1_D10N_, the 18SE level was elevated in excess of that observed upon NOB1 depletion, indicating that the catalytically inactive protein blocks pre-SSU maturation. Expression of FLAG-NOB1_D10N_ also did not restore the levels of the 43S, 30S or 26S pre-rRNAs to those observed in wild-type cells or those expressing FLAG-NOB1_WT_ (**Fig. 3F, G; Supplementary Fig. S3D**), indicating that these defects arise due to lack of NOB1-mediated cleavage of the 3′ end of the 18S rRNA. These results therefore suggest a feedback loop connecting the final maturation step of the 18S rRNA with early pre-rRNA processing steps.

To determine whether the NOB1 phosphorylations at S184 and S201 influence its functions during SSU biogenesis, pre-rRNA levels in cells depleted of endogenous NOB1 and re-expressing wild-type FLAG-NOB1 or FLAG-NOB1_S184A,S201A_ (**Fig. 3H; Supplementary Fig. 3E**) were analysed by northern blotting. As previously (**Fig. 3B-G; Supplementary Fig. S3**), depletion of NOB1 caused accumulation of the 18SE pre-rRNA as well as the 43S and 26S species, concurrent with a reduced 30S, and these pre-rRNA processing defects were rescued by re-expression of wild-type FLAG-NOB1 (**Fig. 3I, J; Supplementary Fig. S3F**). Expression of FLAG-NOB1_S184A,S201A_ restored 30S, 43S and 26S levels similar to expression of the wild-type protein, indicating that phosphorylation of these sites does not influence processing of the 5′ ETS. However, compared to cells expressing wild-type FLAG-NOB1, cells expressing FLAG-NOB1_S184A,S201A_ displayed mildly elevated 21S and reduced 18SE pre-rRNA levels (**Fig. 3I, J; Supplementary Fig. S3F**), suggesting that phosphorylation of these sites contributes to the efficiency of SSU maturation events in ITS1 directly upstream of NOB1-mediated pre-rRNA cleavage.

In summary, these results contribute to the knowledge on post-translational modification of ribosome assembly factors by confirming that NOB1 is a phosphoprotein. S184 and S201 are identified as key phosphorylation sites that are stoichiometrically modified under normal growth conditions in HeLa cells. Although equivalent residues to pS184 and pS201 are aspartates/glutamates in other species suggesting a conserved importance of acidic amino acids at these positions, lack of these phosphorylations does not influence NOB1 association with pre-SSU complexes and only minimally perturbs pre-rRNA processing. Consistent with the location of the phosphorylation sites outside the PIN domain of NOB1, the mild accumulation of the 21S pre-rRNA observed in cells expressing phospho-null NOB1 suggests that these phosphorylations do not affect the catalytic activity of NOB1, but rather subtly influence its function within pre-SSU complexes directly upstream of site 3 cleavage. The nominal effect of dephosphorylation of pS184 and pS201 may reflect their strongly acidic context but a construct lacking this flexible loop was expressed below the endogenous NOB1 level in cells (data not shown), preventing exploration of the functional importance of this feature for ribosome biogenesis.

In the course of this study, it was observed that lack of NOB1 or its catalytic activity change kinetics of 5′ ETS processing. NOB1 has currently only been visualised in pre-SSU particles downstream of 5′ ETS removal (Vanden Broeck and Klinge 2024) so the data presented here indicate the existence of a feedback loop connecting 18S 3′ end maturation to earlier, nucleolar pre-SSU complexes. It is likely that this regulatory circuit involves the recycling of AFs required for early biogenesis steps. The effects on 5′ ETS processing observed when NOB1 is depleted (accumulation of 43S and 26S pre-rRNAs) are also detected in cells lacking the partner of NOB1, PNO1 (Nieto et al. 2020; Sloan et al. 2019). Although accumulation of the 26S pre-rRNA is observed when pre-SSU export to the cytosol is perturbed (Morello et al. 2011), PNO1 is required for formation of early pre-SSU particles in the nucleolus, rather than being involved at the export stage (Nieto et al. 2020). Given the similarity of the early pre-rRNA processing defects observed upon their depletion, it is tempting to speculate that the feedback loop connecting 18S 3′ end maturation to 5′ ETS processing may involve both proteins.

## MATERIALS AND METHODS

### Molecular cloning

The coding sequences (CDS) of NOB1 (*Hs*NOB1: NM_014062.3, *Mm*NOB1: NM_026277.3) were amplified by polymerase chain reaction (PCR) from complementary DNA (cDNA) prepared from HEK293 Flp-In T-REx cells (Thermo Fisher Scientific) or mouse total RNA, respectively. The plasmids pcDNA5-2xFLAG-His_6_ and pGEX-4T-1 were used as vectors for the expression of proteins in human cells and the recombinant expression of proteins in *E. coli*, respectively (**Supplementary Table S2**). Cloning was performed using Gibson assembly in combination with site-directed mutagenesis using primers listed in **Supplementary Table S3**. The presence of mutations was confirmed by Sanger sequencing.

### Cell culture and generation of stably transfected cell lines

HeLa Flp-In and mouse C2C12 cells were cultured at 37°C with 5% CO_2_ in Dulbecco’s modified Eagle’s medium (DMEM; Thermo Fisher Scientific) supplemented with 10% foetal bovine serum (FBS; Sigma-Aldrich) and 1% penicillin–streptomycin (Gibco). Stably transfected HeLa Flp-In cell lines were generated by transfection with pcDNA5-based plasmids encoding N-terminally 2xFLAG-His_6_ (FLAG)-tagged proteins and the pOG44 plasmid encoding a Flp recombinase using XtremeGENE9 (Roche) transfection reagent according to the manufacturer’s instructions. Cells were selected for appropriate integrations in the Flp-In locus using 200 µg/mL of Hygromycin B and 1 µg/mL of Blasticidin S. Expression of the FLAG-tagged proteins was induced by addition of 1 µg/mL tetracycline for 24 h before harvesting. Transiently transfected HeLa Flp-In or C2C12 cells were generated by transfection with pcDNA5-based plasmids encoding FLAG-tagged proteins (**Supplementary Table S2**) using Lipofectamine 2000 (Thermo Fisher Scientific) according to the manufacturer’s instructions and harvested 24 h after transfection. Protein expression was induced with tetracycline at the time of transfection.

### RNAi-mediated depletion

HeLa Flp-In cells were transfected with 50 nM siRNAs (**Supplementary Table S4**) using Lipofectamine RNAiMAX (Thermo Fisher Scientific) according to the manufacturer’s instructions. Cells were harvested 72 h after transfection. For experiments combining RNAi with expression of transgenes from the Flp-In locus, cells were transfected with appropriate siRNAs as above and treated with 1 µg/mL tetracycline 24 h prior to harvesting.

### Northern blotting of pre-rRNAs

Total RNA was extracted from HeLa cells using TRI-reagent (Sigma-Aldrich) according to the manufacturer’s instructions. 2-4 µg total RNA were mixed with 5 volumes glyoxal loading dye (61% dimethylsulfoxide (DMSO), 20% glyoxal, 5% glycerol, 10 mM PIPES, 30 mM Bis-Tris, 1 mM ethylenediaminetetraacetic acid (EDTA)) and denatured at 55°C for 1 h. RNAs were separated at 60 V for 16 h in a 1.2% agarose-BPTE (Bis-Tris-PIPES-EDTA; 10 mM PIPES, 30 mM Bis-Tris, 1 mM EDTA) gel, then hydrolysed *in situ* with 100 mM NaOH for 20 min and washed 2x 15 min with Tris-salt buffer (0.5 M Tris-HCl pH 7.4, 1.5 M NaCl) and 1x 15 min with 6x SSC buffer (saline-sodium-citrate; 90 mM tri-sodium citrate, 300 mM NaCl) before vacuum transfer (300 mbar, 2 h) to a Hybond-N+ (Cytiva) nylon membrane. RNAs were crosslinked to the membrane with 2x 120 mJ/cm^2^ at 254 nm wavelength. Northern blot oligonucleotide probes (5′ ITS1: 5′-CCTCGCCCTCCGGGCTCCGTTAATGATC-3′; mt-16S: 5′-TTCTATAGGGTGATAGATTGGTCC-3′) were labelled at the 5’ end with γ-[^32^P] using T4 polynucleotide kinase (PNK, Thermo Fisher Scientific). Membranes were pre-hybridised with SES1 buffer (250 mM NaPi pH 7.4, 7% sodium dodecyl sulfate (SDS), 1 mM EDTA pH 8) for 30 min before overnight hybridisation with radioactively-labelled probes (1 µM oligo, 1x PNK buffer A (Thermo Fisher Scientific), 20 µCi γ-[^32^P]-ATP, 0.5 U/µL PNK incubated at 37°C for 40 min in a 20 µL reaction volume) diluted in SES1 at 37°C. Membranes were washed with 6x SSC followed by washing with 2x SSC + 0.1% SDS for 30 min each at 37°C. Air-dried membranes were exposed to phosphorimager screens and images were captured using an Amersham TYPHOON (Cytiva). Images were processed using Image Studio software (LICORbio), quantified using ImageQuant TL 11.0 (Cytiva), and heatmaps and bar graphs were made with GraphPad Prism (v.10.4.2).

### Preparation of PI/AP-treated cell extracts and total protein extraction

Cell pellets were resuspended in RIPA buffer (150 mM NaCl, 1% NP-40, 50 mM Tris-HCl pH 7.4, 1.5 mM MgCl_2_, 0.5% sodium deoxycholate, 0.1% SDS) supplemented with cOmplete EDTA-free protease inhibitor cocktail tablets (Roche) and, where indicated, either PhosSTOP phosphatase inhibitor tablets (PI, Roche) or alkaline phosphatase (FastAP, Thermo Fisher Scientific) and incubated at 4°C (PI-treated) or 37°C (AP-treated) for 45 min. Lysates were centrifuged at 956 xg, 4°C for 5 min after which proteins from the supernatant were precipitated with 12.5% trichloroacetic acid (TCA) and centrifugation at 956 xg, 4°C for 5 min. Protein pellets were washed twice with cold acetone, centrifuging at 17,949 xg, 4°C for 5 min after each wash, dried and resuspended in 1x NuPAGE loading dye containing 50 mM DTT, supplemented with 10 mM Tris-HCl pH 8.8. Protein concentrations were determined from cell lysate in RIPA buffer using the Pierce BCA Protein Assay kit (Thermo Fisher Scientific) according to the manufacturer’s instructions.

### Bis-Tris and Phos-tag gel electrophoresis and western blotting

10 µg total protein were heated at 70°C for 10 min and separated by electrophoresis in 8-10% Bis-Tris-polyacrylamide gels in Tris-MOPS running buffer (50 mM Tris, 50 mM MOPS, 0.1% SDS), supplemented with 4.8 mM sodium bisulfite, and 200 mM Tris-HCl pH 9 anode buffer at 40 mA per gel with maximum 90 V. Phos-tag gels were prepared as Bis-Tris-polyacrylamide gels containing 50 µM Phos-tag solution (FUJIFILM Wako Pure Chemical Corporation) and 100 µM ZnCl_2_ in both the stacking and resolving gels. Proteins were transferred onto PVDF membranes (Sigma-Aldrich, Bio-Rad) by wet-transfer at 60 V for 90 min. Membranes were blocked with 5% w/v milk in TBS-T (Tris-buffered saline supplemented with 0.1% (v/v) Tween20) for 1 h at RT before incubation with primary antibodies (**Supplementary Table S5**) diluted in 5% (w/v) milk in TBS-T at 4°C o/n or 2 h at RT. Membranes were washed with TBS-T, incubated with secondary antibodies (**Supplementary Table S5**) in TBS-T supplemented with 0.01% SDS for 1 h at RT and again washed with TBS-T. Membranes were then scanned using an Odyssey CLx Imager scanner (LICORbio) and images processed using Image Studio software (LICORbio).

### Anti-FLAG immunoprecipitation (IP) for phosphoprotein gel staining and phosphorylation mass spectrometry

Cell pellets from 2x 15 cm diameter dishes at 80-90% confluence were resuspended in 1 mL IP buffer (50 mM Tris-HCl pH 7.4, 250 mM NaCl, 0.5 mM EDTA, 0.5% Triton-X-100, 10% glycerol, cOmplete EDTA-free protease inhibitor cocktail tablets (Roche), PhosSTOP phosphatase inhibitor tablets (Roche)) and sonicated at 20% amplitude for 3x 15 s, 0.3 s on/0.7 s off with 20 s intervals. Lysed cells were centrifuged at 20,000 xg, 4°C for 10 min and 0.2% of the cleared lysate reserved as input. Anti-FLAG M2 magnetic beads (Sigma-Aldrich; 30 µL slurry per sample) were equilibrated in 3x 500 µL IP buffer. The cleared lysate was then added to the beads and incubated at 4°C for 1 h, rotating “head-over-tail”. The lysate was removed from the beads, the beads washed 5x with 1 mL of a stringent wash buffer (50 mM Tris-HCl pH 7.4, 1 M NaCl, 5 mM EDTA, 1% Triton-X-100, 10% glycerol, 2 M urea, 0.1% SDS, Complete EDTA-free protease inhibitor cocktail tablets (Roche), PhosSTOP phosphatase inhibitor tablets (Roche)). The beads were then resuspended in 250 µL IP buffer and transferred to a 0.5 mL tube. The buffer was removed again, 150 µg/ml 3xFLAG peptide in IP buffer was added and incubated at 4°C for 30 min, rotating “head-over-tail”. The eluate was transferred to a fresh tube. Proteins from the input and eluate were precipitated by addition of 21% TCA, vortexing and incubation on ice for 20 min. Samples were then centrifuged at 20,000 xg, 4°C for 20 min, the supernatant removed and protein pellets washed 5x with 500 µL cold acetone. Air-dried pellets were resuspended in 30 µL NuPAGE loading dye (1x NuPAGE with 50 mM DTT) supplemented with 10 mM Tris-HCl pH 8.8. Proteins in loading dye were separated by Bis-Tris gel electrophoresis, next to PeppermintStick phosphoprotein molecular weight standard (Molecular Probes, used according to the manufacturer’s instructions), followed by staining with Pro-Q Diamond phosphoprotein gel stain (Invitrogen) according to the manufacturer’s instructions. Phosphorylated proteins were visualised using a Typhoon 9400 (Amersham) with excitation at 532 nm and 560 nm longpass emission filter. Alternatively, samples were used for phosphorylation mass spectrometry.

### Phosphorylation mass spectrometry

Protein bands corresponding to the FLAG-NOB1 protein were excised from the NuPAGE gel. Proteins were reduced with dithiothreitol, alkylated with iodoacetamide and in-gel digested overnight by trypsin. Extracted peptides were dried in a vacuum concentrator.

In order to enrich phosphopeptides, dry peptide pellets were dissolved in 60 μL loading buffer (5% glycerol, 80% acetonitrile (ACN), 5% trifluoroacetic acid (TFA)) and loaded onto self-packed TiO_2_ tips (200 µL pipette tips with Titansphere Phos-TiO 5 µm beads, MZ-Analysentechnik GmbH, 5020-75000, bed height 3 mm), which were pre-equilibrated with 60 μL wash buffer I (80% ACN, 5% TFA) followed by 60 μL loading buffer. The TiO_2_ tips were then washed 3x with 60 µL loading buffer, three times with 60 µL wash buffer I and once with 60 µL 60% ACN, 0.1% TFA. Peptides were eluted from the tips by three consecutive additions of 40 μL 10.3 N NH_4_OH (pH ≥ 10.5), and immediately acidified with formic acid (FA, final concentration approx. 1.4%, pH ≤ 3). Eluted phosphopeptides were dried in the vacuum concentrator, dissolved in 20 µL 2% ACN, 0.1% TFA and injected into a nanoflow ultra-high pressure liquid chromatography UltiMate 3000 UHPLC system coupled to an Orbitrap Exploris 480 mass spectrometer equipped with an electro-spray ionization (ESI) Nanospray Flex Ion source (Thermo Fisher Scientific) operating in positive mode. The peptides were pre-concentrated and desalted on a trap column (PepMap Neo C18, 5 μm, 0.3 × 5 mm, Thermo Fisher Scientific) and then separated on a 27 cm self-packed main column (inner diameter of 75 µm, Reprosil-Pur 120 C18-AQ 3 μm beads, Dr. Maisch GmbH). The data were acquired in data-dependent acquisition (DDA) mode using a 58-minute method with a 10 to 40% gradient of solvent B (80% acetonitrile, 0.1% FA in water) at a flow rate of 300 nL/min. Cycle time was set to 3 s. The resolution settings were 120,000 and 30,000 full width, half maximum (FWHM) for the MS1 and MS2 scans, respectively. MS1 scan range was 350-1550 *m/z*. MS2 maximum injection time was 60 ms. MS1 and MS2 normalised AGC targets were 300 and 100%, respectively. MS2 Fragmentation was achieved with higher-energy collisional dissociation (HCD) and a normalised collision energy setting of 28%.

MS raw files were searched with MASCOT v.2.3.02 (Matrix Science) against the UniprotKB human reference proteome (release 2024-07-24). Trypsin was set as protease, 2 missed cleavages were allowed. Carbamidomethylation of cysteine (fixed modification) as well as oxidation of methionine, phosphorylation of serine, threonine and tyrosine (variable modifications) were considered. The MASCOT search results were loaded into Scaffold v.5.3.3 (Proteome Software Inc.). The results were filtered to protein and peptide thresholds of 99 and 95%, respectively.

### Sucrose density gradient centrifugation

Cell pellets from 1x 15 cm diameter dishes at 80-90% confluence were resuspended in 500 µL lysis buffer (50 mM Tris-HCl pH 7.5, 100 mM NaCl, 5 mM MgCl_2_, 1 mM DTT, cOmplete EDTA-free protease inhibitor cocktail tablets (Roche), PhosSTOP phosphatase inhibitor tablets (Roche)) and sonicated at 20% amplitude, 3x 15 s, 0.3 s on/0.7 s off with 20 s intervals. The lysate was centrifuged at 20,000 xg, 4°C for 10 min and the cleared lysate transferred to a new tube. 10-45% sucrose density gradients were prepared by layering 10% and 45% sucrose solutions (50 mM Tris-HCl pH 7.5, 100 mM NaCl, 5 mM MgCl_2_, 1 mM DTT, 10/45% sucrose) on top of each other in rotor tubes and mixing using a Gradient Station *ip* (Biocomp) (rotate for 1:25 min at 82.0° and speed 19). The prepared gradients were cooled at 4°C for 1 h before replacing 400 µL from the top with the cleared lysate. 10% of the lysate were reserved as input. Complexes were separated on the density gradients by centrifugation at 23,500 rpm, 4°C for 16 h in an Optima XPN-80 ultracentrifuge (Beckman Coulter) with SW 40 Ti rotor (Beckman Coulter). Fractionation of the samples was performed with a Gradient Station *ip* (Biocomp) and connected fraction collector (Gilson), with continuous monitoring of RNA content through measuring absorbance at 254 nm. Proteins were precipitated with 12.5% TCA and incubation on ice for 20 min before centrifugation at 20,000 xg, 4°C for 20 min. Pellets were washed with cold acetone, dried and resuspended in 100 µL NuPAGE loading dye supplemented with 10 mM Tris-HCl pH 8.8 for analysis by Bis-Tris gel electrophoresis and western blotting. 10% gels were run as described above at 25 mA until samples reached the separating gel, then 30 mA for 2 h total. Transfer, blocking and incubation with antibodies was carried out as described above. Alternatively, relevant fractions were pooled and proteins precipitated for analysis by Phos-tag gel electrophoresis and western blotting.

### Recombinant protein expression in E. coli and purification

Relevant plasmids were used to transform *E. coli* BL21 CodonPlus, which were selected on LB supplemented with ampicillin (Amp; 100 µg/mL) and chloramphenicol (Chl; 34 µg/mL). Single colonies were used to inoculate 100 mL pre-cultures in LB Amp/Chl and grown at 37°C o/n. 1 L cultures were then inoculated with 20 mL pre-culture and grown to OD_600_ ≈ 0.6-0.8 at 37°C. Cultures were cooled rapidly, induced with 1 mM IPTG and grown at 18°C o/n. Pre- and post-induction samples were taken. Cells were pelleted at 6,000 xg, 4°C for 10 min, washed with PBS and again pelleted at 4,000 xg, 4°C for 10 min. Pellets were resuspended in 20 mL lysis buffer (20 mM Tris-HCl pH 7.4, 300 mM KCl, 5 mM MgCl_2_, 10% glycerol, 0.1% Tween20, cOmplete EDTA-free protease inhibitor cocktail tablets (Roche)) and sonicated on ice for 2x 3 min at 70% amplitude, 0.5 s on/0.5 s off. The lysate was cleared by centrifugation at 20,000 xg, 4°C for 30 min and samples from the pellet and lysate were taken. Glutathione Sepharose Fast Flow beads (Cytiva, 1 mL slurry) were equilibrated in 3x 5 mL lysis buffer before the cleared lysate was added to the beads and incubated at 4°C for 2 h while slowly rotating. Beads with bound protein were sedimented by centrifugation at 1,000 xg for 5 min, a sample from the supernatant taken as “flow-through” and the beads washed with 2x 50 mL lysis buffer. The beads were then resuspended in 2 mL lysis buffer containing 50 mM glutathione pH 8.0 and incubated on a rotating wheel at 4°C for 1 h to elute the protein. The supernatant was collected using Mobicol columns (MoBiTec) and dialysed against dialysis buffer (20 mM Tris-HCl pH 7.4, 300 mM KCl, 5 mM MgCl_2_, 20% glycerol) at 4°C o/n, followed by centrifugation at 20,000 xg, 4°C for 20 min. Samples from the supernatant before and after dialysis were taken as eluate and final, respectively. Samples were separated by Bis-Tris gel electrophoresis and detected by Coomassie staining.

### Phosphorylation assay

10 µg recombinant GST-NOB1_WT_/GST-NOB1_S184A,S201A_ were mixed with 1 µL casein kinase II (2,000 U/µL, New England Biolabs), 2 µCi γ-[^32^P]-ATP, 0.1 mM ATP and 1x protein kinase buffer (New England Biolabs) in a 10 µL reaction and incubated at 30°C for 20 min. The reaction products were separated by denaturing Bis-Tris gel electrophoresis as described above. The gel was dried and exposed to a phosphorimager screen overnight and images obtained with an Amersham TYPHOON (Cytiva).

### Multiple Sequence Alignment

Multiple sequence alignment was performed with Clustal Omega and manually curated. The following species were included: *Homo* sapiens (*Hs*NOB1, Q9ULX3), *Mus musculus* (*Mm*NOB1, Q8BW10), *Danio rerio* (*Dr*NOB1, I3ISV3), *Drosophila melanogaster* (*Dm*NOB1, Q9W2Y4), *Caenorhabditis elegans* (*Ce*NOB1, Q9N3D3), *Candida albicans* (*Ca*NOB1, Q59SR4), *Saccharomyces cerevisiae* (*Sc*NOB1, Q08444), *Schizosaccharomyces pombe* (*Sp*NOB1, Q9UTK0).

## DATA AVAILABILITY

The mass spectrometry data generated for this study have been deposited at the ProteomeXchange Consortium via the MassIVE partner repository [<u>https://massive.ucsd.edu</u>] under the following accession codes: MSV000101496 | <u>PXD077313</u>.

## ACKNOWLEDGEMENTS

We thank Prof. Thomas Mayer (University of Konstanz) for sharing the HeLa Flp-In cell line, Prof. Ralph Kehlenbach for sharing the mouse C2C12 cell line and the lab of Prof. Peter Rehling for mouse total RNA. We thank Monika Raabe for assistance with phosphorylation mass spectrometry and Sabrina Stille for input on recombinant protein purification. This work was funded by the Deutsche Forschungsgemeinschaft (DFG, German Research Foundation), project number 469281184, SFB1565 to HU (P04), MTB (P06), KEB (P12), SL (P17) and CL (Z02).

## SUPPLEMENTARY INFORMATION

**Supplementary Figure S1.**
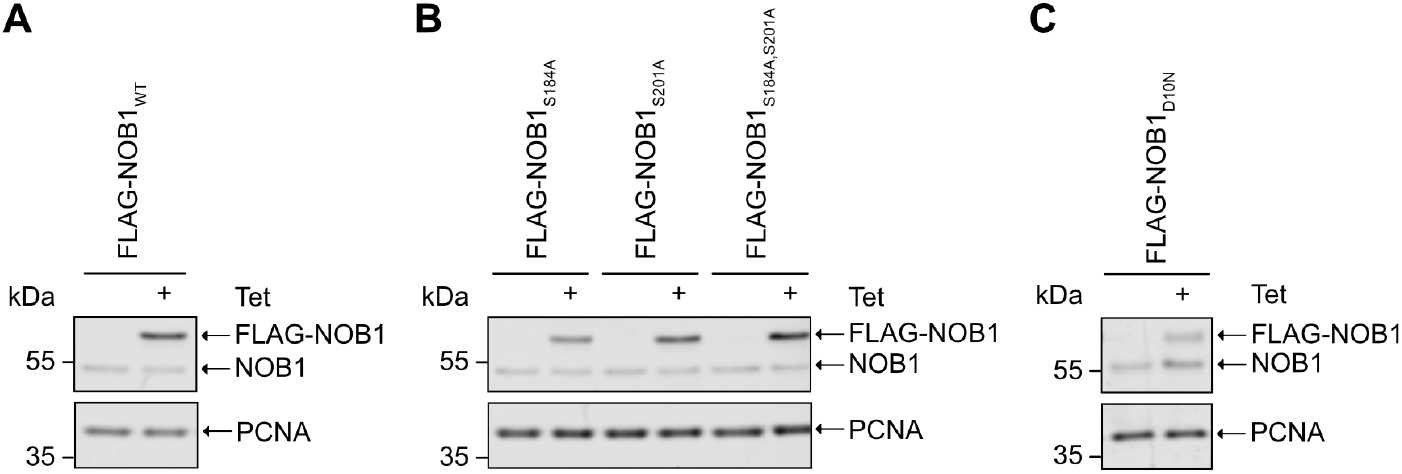
Expression of NOB1 from stable cell lines. **(A-C)** Western blot analysis (anti-NOB1) of stably transfected HeLa cells expressing FLAG-NOB1_WT_, FLAG-NOB1_S184A_, FLAG-NOB1_S201A_, FLAG-NOB1_S184A,S201A_ or FLAG-NOB1_D10N_ from a tetracycline (Tet)-inducible promoter. PCNA served as a loading control.

**Supplementary Figure S2.**
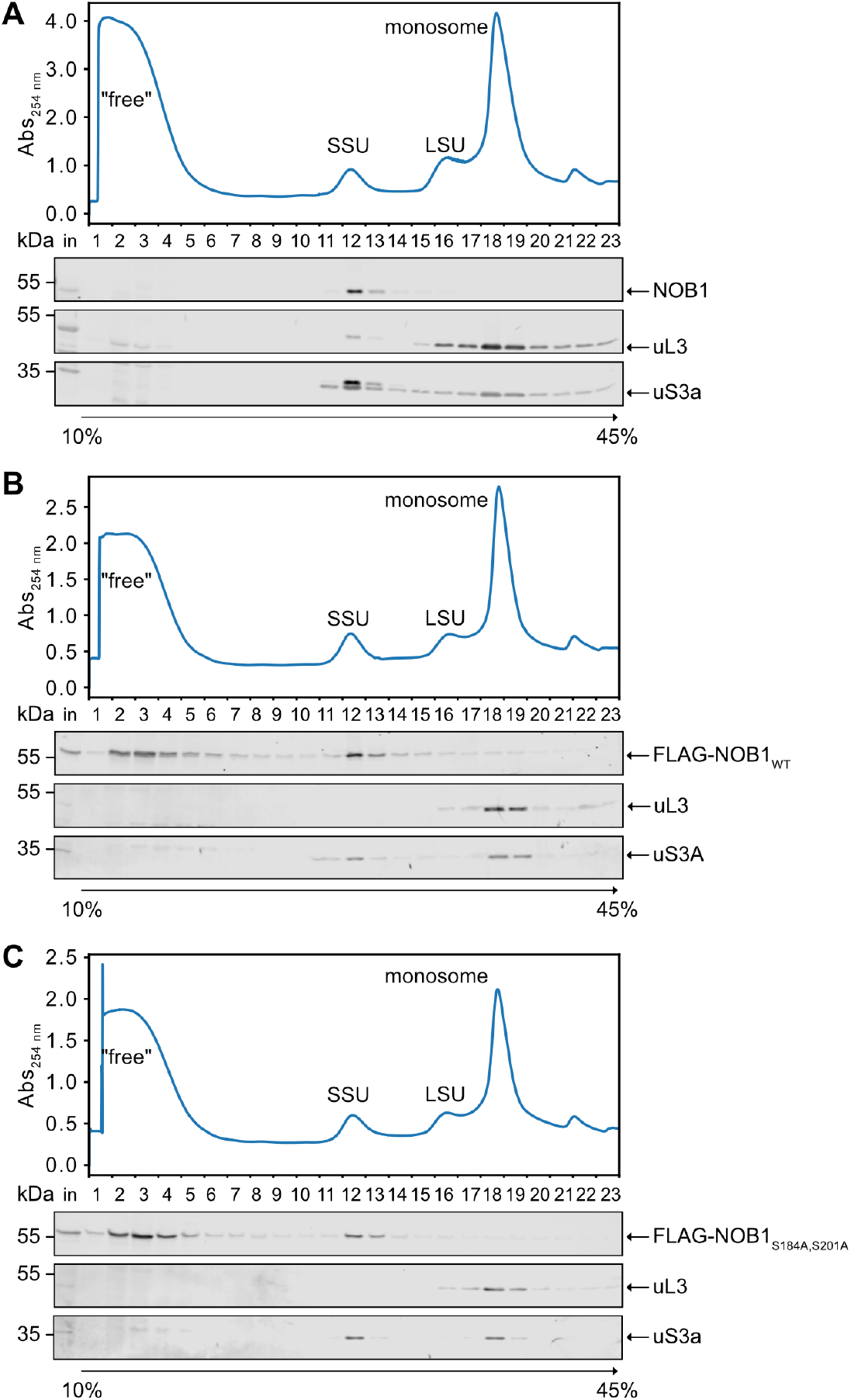
NOB1, FLAG-NOB1_WT_ and phospho-null FLAG-NOB1 association with pre-ribosomes. **(A-C)** Phosphatase inhibitor-treated extracts from HeLa cells (A) and HeLa cell lines expressing FLAG-NOB1_WT_ (B) or FLAG-NOB1_S184A,S201A_ (C) were separated by sucrose density gradients. The absorbance profiles at 254 nm are shown with peaks corresponding to (pre-)SSU, (pre-)LSU and monosomes/90S pre-ribosomes indicated. Proteins in each fraction of the gradients were analysed by western blotting using anti-NOB1 (A), anti-FLAG (B, C), and uL3 and uS3a antibodies. Some data are also shown in **Fig. 2E**.

**Supplementary Figure S3.**
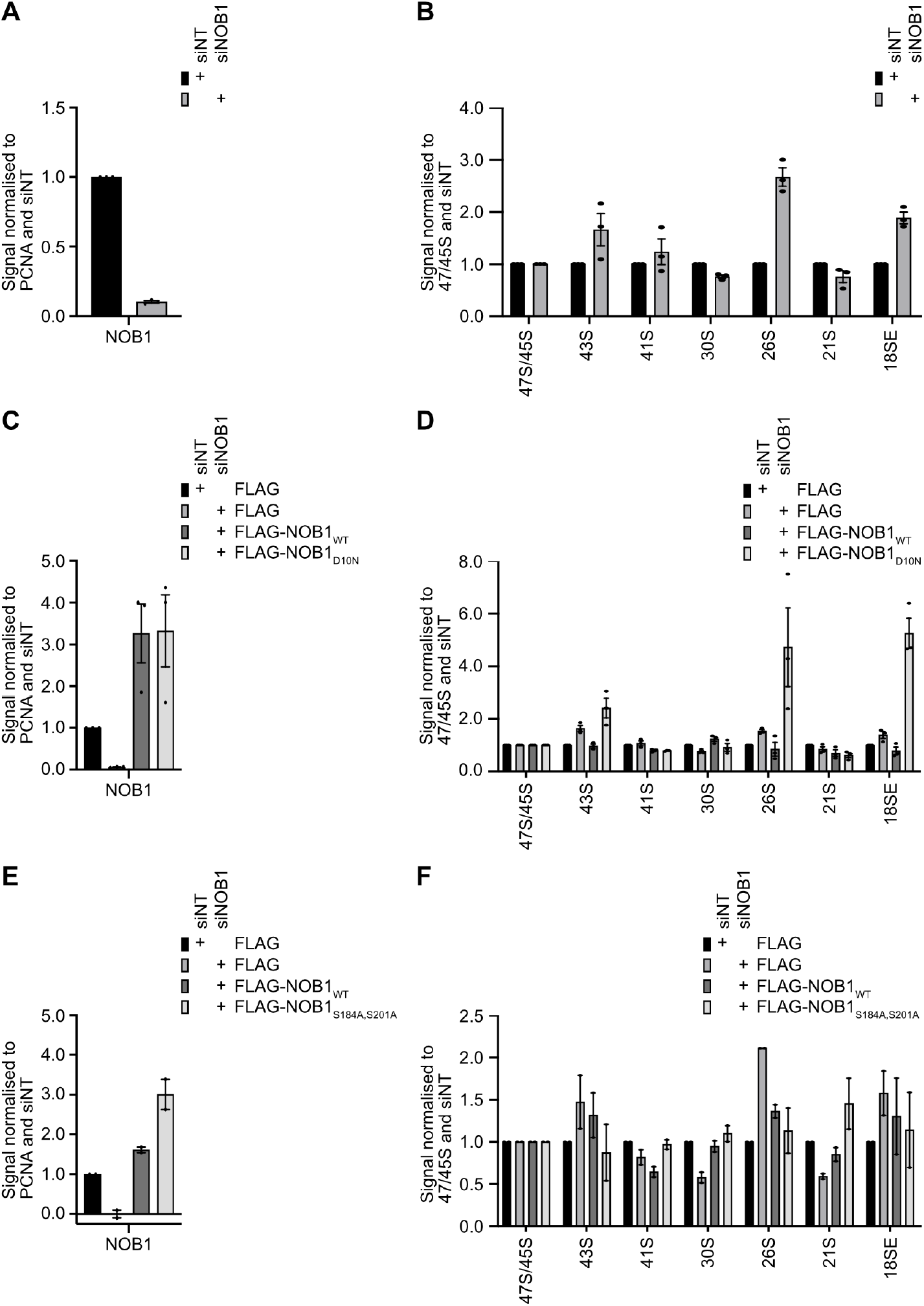
NOB1 expression levels and pre-rRNA quantifications. **(A)** Quantification of NOB1 knockdown efficiency after treatment with non-target (siNT) or NOB1 (siNOB1) siRNAs (Fig. 3B). (B) Quantification of pre-rRNA intermediates from (**Fig. 3C**). **(C)** Quantification of NOB1 expression levels after siRNA-mediated knockdown and expression of a FLAG-tagged version of NOB1 (**Fig. 3E**). **(D)** Quantification of pre-rRNA intermediates from (**Fig. 3F**). **(E)** Quantification of NOB1 expression levels after siRNA-mediated knockdown and expression of a FLAG-tagged version of NOB1 (**Fig. 3H**). **(F)** Quantification of pre-rRNA intermediates from (**Fig. 3I**). All data are presented as mean of three (A-D) or two (E-F) independent experiments, normalised to PCNA (A, C, E) or 47S/45S (B, D, F) and scaled to siNT, with error bars representing s.e.m. (standard error of the mean) and individual data points shown.

**Supplementary Table S1.**
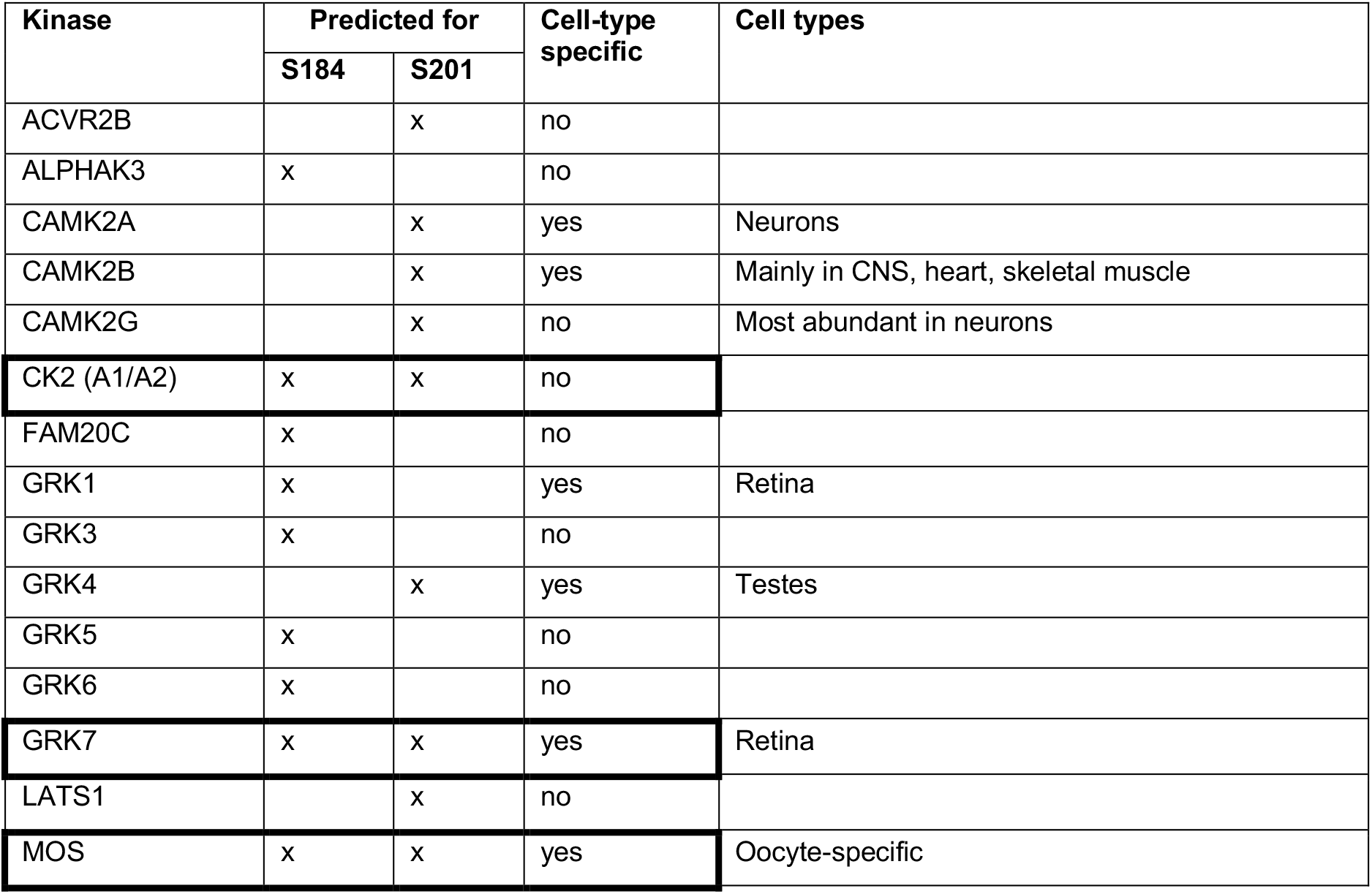
Kinase predictions for NOB1 S184 and S201. The top ten kinases predicted to phosphorylate S184 or S201 according to (Johnson et al. 2023) are listed with information on whether expression of the kinase is cell-type specific (according to the Human Protein Atlas (Uhlén et al. 2015) or UniProt (The UniProt Consortium 2025)). As S184 and S201 are in close vicinity to each other and both highly phosphorylated, kinases predicted to target both serines are highlighted with boxes.

**Supplementary Table S2.** Plasmids used for expression of NOB1 and its derivatives in human and bacterial cells.

| Internal ref # | Name | Purpose |
| --- | --- | --- |
| pMB-064 | pcDNA5-FLAG | Generation of stable cell line |
| pMB-2132 | pcDNA5-FLAG-NOB1 WT | Generation of stable cell line |
| pMB-2133 | pcDNA5-FLAG-NOB1 S184A | Generation of stable cell line |
| pMB-2134 | pcDNA5-FLAG-NOB1 S201A | Generation of stable cell line |
| pMB-2136 | pcDNA5-FLAG-NOB1 S184A S201A | Generation of stable cell line |
| pMB-2448 | pcDNA5-FLAG-NOB1 D10N | Generation of stable cell line |
| pMB-2449 | pcDNA5-FLAG-NOB1 S325A | Transient transfection |
| pMB-2135 | pcDNA5-FLAG-NOB1 S352A | Transient transfection |
| pMB-2450 | pcDNA5-FLAG-*Mm*NOB1 WT | Transient transfection |
| pMB-2451 | pcDNA5-FLAG-*Mm*NOB1 S192A | Transient transfection |
| pMB-2452 | pGEX-4T-1 NOB1 WT | Recombinant expression |
| pMB-2453 | pGEX-4T-1 NOB1 S184A S201A | Recombinant expression |

**Supplementary Table S3.**
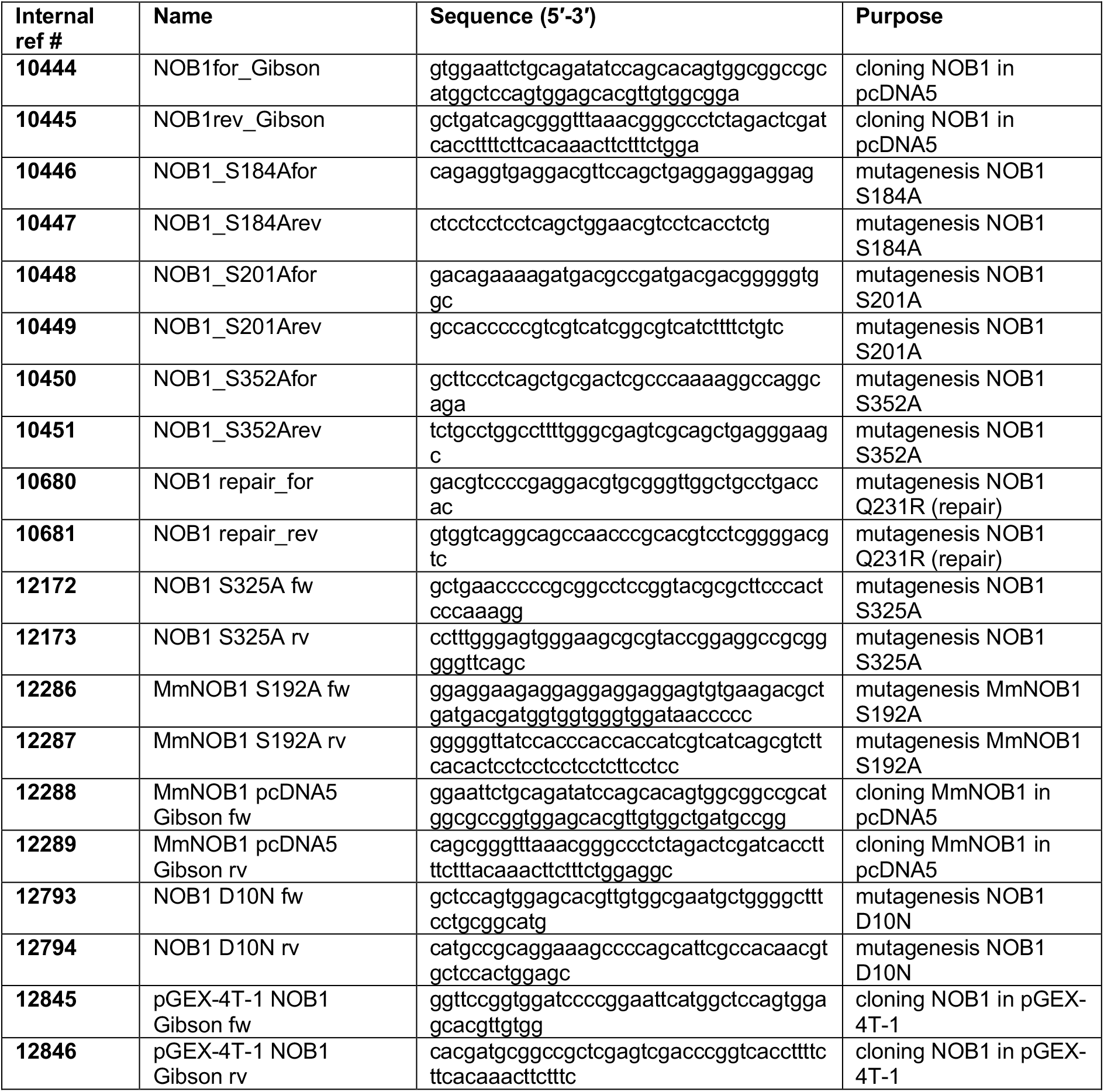
Oligonucleotide primers used for molecular cloning and site-directed mutagenesis.

**Supplementary Table S4.**
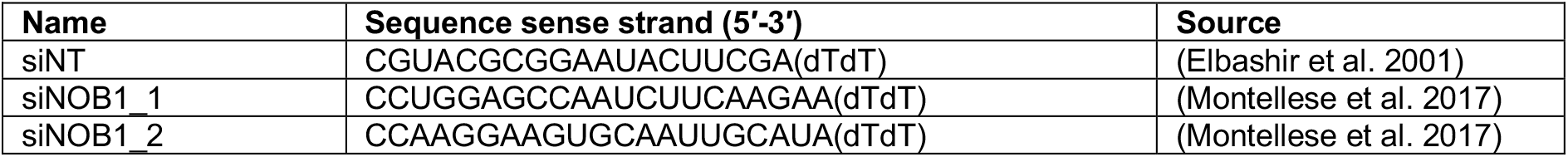
siRNAs used for RNAi-mediated depletion.

**Supplementary Table S5.** Antibodies and dilution factors.

| Antibody | Host species | Dilution | Source | Catalogue # |
| --- | --- | --- | --- | --- |
| anti-FLAG | rabbit | 1:2000 | Sigma-Aldrich | F7425 |
| anti-FLAG | mouse | 1:5000 | Sigma-Aldrich | F3165 |
| anti-NOB1 | rabbit | 1:1000 | Proteintech | 10091-2-AP |
| anti-PCNA | mouse IgG2a | 1:100 | Santa Cruz | sc-56 |
| anti-uL3 | rabbit | 1:5000 | Proteintech | 11005-1-AP |
| anti-uS3a | rabbit | 1:5000 | Proteintech | 14123-1-AP |
| IRDye 800CW anti-rabbit IgG | goat | 1:10000 | LI-COR | 926-32211 |
| IRDye 680LT anti-mouse IgG2a | goat | 1:10000 | LI-COR | 926-68051 |
| IRDye 800CW anti-mouse IgG | donkey | 1:10000 | LI-COR | 926-32212 |
| IRDye 680LT anti-rabbit IgG | donkey | 1:10000 | LI-COR | 926-68023 |

